# Connexin43 drives mitochondrial biogenesis and organelle stress in BRAF-mutant melanoma cells

**DOI:** 10.64898/2026.11.18.767001

**Authors:** Amanda Guitián-Caamaño, Eva Fernández-Bejarano, Cristina De Jesús-Sen, Diego González-Pérez, Paula Carpintero-Fernández, Teresa Calleja-Chuclá, Carlos Fernandez-Lozano, Eduardo Fonseca, Aurora Gomez-Duran, María D. Mayán

**Affiliations:** CellCOM Research Group. Center for Research in Nanomaterials and Biomedicine (CINBIO) and Institute of Biomedical Research of Ourense-Pontevedra-Vigo (IBI), University of Vigo. The Galician Health Service, SERGAS. Olimpia Valencia building, Campus Universitario Lagoas Marcosende, 36310. Vigo, Spain. Before: CellCOM group. INIBIC, UDC, SERGAS. A Coruña. Spain; Conway Institute of Biomolecular and Biomedical Research. School of Mechanical and Materials Engineering, Engineering and Materials Science Centre. University College Dublin, Dublin D04 V1W8, Ireland; Pharmacy Service CHUAC.INIBIC, Servizo Galego de Saúde (SERGAS), Universidade da Coruña (UDC), A Coruña, Spain; MitoPhenomics Lab. Centro Singular de investigación en Medicina Molecular y Enfermedades Crónicas (CiMUS), Biochemistry Department. University of Santiago de Compostela (USC). Santiago de Compostela, 15782. A Coruña, Spain; Genetics and Systems Biology Department, Foundation Health Research Institute of Santiago de Compostela (IDIS), 15706, Santiago de Compostela, A Coruña, Spain; Department of Computer Science and Information Technologies, Faculty of Computer Science, Universidade da Coruña, A Coruña, 15071, Spain

**Author notes:** **Corresponding authors: María D. Mayán.** CellCOM Research Group. Center for Research in Nanomaterials and Biomedicine (CINBIO) and Institute of Biomedical Research of Ourense-Pontevedra-Vigo (IBI), University of Vigo. The Galician Health Service, SERGAS. Olimpia Valencia building, Campus Universitario Lagoas Marcosende, 36310, Pontevedra, Spain., Aurora Gomez-Duran. MitoPhenomics Lab. Centro Singular de investigación en Medicina Molecular y Enfermedades Crónicas (CiMUS), and Foundation Health Research Institute of Santiago de Compostela (IDIS). Biochemistry Department. University of Santiago de Compostela (USC). Santiago de Compostela, 15782. A Coruña, Spain. These authors contributed equally to this work (Amanda Guitián-Caamaño, Eva Fernández-Bejarano).

**Keywords:** BRAF mutation, Connexins, Cx43, mitochondria, BRAF and MEK inhibitors

## Abstract

Metabolic flexibility helps cancer cells survive targeted therapy, and BRAF-mutant melanoma rewires mitochondrial metabolism in response to BRAF/MEK inhibitors (BRAF/MEKi). We previously showed that connexin43 (Cx43) acts as a tumour suppressor in BRAF-mutant melanoma and enhances the response to BRAF/MEKi. Here, we investigated whether mitochondrial remodelling contributes to these effects. Cx43 upregulation promoted mitochondrial biogenesis including selective increase of CI and CV, enhanced oxidative phosphorylation (OXPHOS), and increased reactive oxygen species (ROS) production, alongside DRP1-associated mitochondrial fragmentation. Despite enhanced mitochondrial biogenesis and OXPHOS, Cx43 upregulation was associated with respiratory chain remodelling, characterised by loss of complex IV, increased ER and mitochondrial stress, and enhanced mTORC1-associated signalling. These changes were accompanied by an adaptive unfolded protein response that did not engage the pro-apoptotic ATF4–CHOP axis. Notably, BRAF/MEK inhibition largely attenuated the mitochondrial biogenesis phenotype and most of the stress markers associated with Cx43 upregulation, while promoting mitochondrial network elongation and markedly increasing mitochondrial ROS. Despite this impact, Cx43-expressing tumour cells retained higher membrane potential, ROS levels and survival under glucose-limiting conditions. Collectively, these findings reveal a dissociation between the mitochondrial effects of Cx43 and its enhancement of the response to BRAF/MEKi, highlighting the complexity of Cx43-mediated metabolic regulation in BRAF-mutant melanoma. By distinguishing mitochondrial remodelling from therapeutic sensitisation, this study provides new insights into the response to BRAF/MEK inhibition and highlights mTORC1-associated signalling and mitochondrial oxidative stress as potential therapeutic vulnerabilities for further investigation.

## Introduction

Mutations in the *BRAF* proto-oncogene are associated with the development and progression of many types of cancer. *BRAF* is a component of the Ras/Raf/MEK/ERK mitogen-activated protein kinase (*MAPK*) signalling pathway, whose main function is to regulate cell growth, migration, and proliferation [1, 2]. The *BRAF*^V600E^ mutation is the most prevalent, leading to constitutive activation, enhancement of survival and uncontrolled cell growth. This mutation has been linked to melanoma, colorectal cancer, and up to 8% of all cancer types [3, 4]. The development of *BRAF* and *MEK* inhibitors (*BRAF/MEKi*) marked a paradigm shift in the treatment of *BRAF*-mutated melanoma, as well as lung or anaplastic thyroid cancers [5]. At the molecular level, more than 50% of melanoma cases harbour a *BRAF* mutation [6]. Melanoma is the most lethal form of skin cancer, and its incidence continues to increase worldwide, representing a significant challenge in cancer treatment. *BRAF/MEK*i, either alone or in combination with immunotherapies, have become the standard therapy for melanoma patients, leading to a significant advancement in its management [7, 8]. Despite their effectiveness, resistance to BRAF/MEKi therapies remains a major clinical challenge.

Reprogramming of energy metabolism is a recognized hallmark of cancer [9]. Cellular metabolism is closely linked to oncogenes and serves as a key mechanism that drives continuous cell growth and proliferation [10]. The *BRAF* oncogene affects metabolism by regulating different signalling pathways [11] and has emerged as a critical regulator of the *Warburg effect*. This metabolic shift causes cancer cells to rely more on glycolysis than mitochondrial oxidative phosphorylation (OXPHOS), promoting their survival and growth [11, 12]. Besides, this metabolic shift is further evidenced by increased glucose uptake in *BRAF*-mutant melanomas compared to normal tissues [12]. Clinically, patients with *BRAF*-mutant melanomas show increased serum lactate, pointing to reduced OXPHOS function [13]. In *in vitro* models, *BRAF*^V600E^ contributes to melanoma cell survival under oxidative metabolic stress conditions through melanocyte master regulator (microphthalmia-associated transcription factor, MITF) and the mitochondrial biogenesis co-activator PGC1α [14–17]. The *BRAF*^V600E^ mutation is also linked to metabolic rewiring [18, 19], where *BRAF*^V600E^ mutant melanomas appear to downregulate the expression of proteins involved in fatty acid oxidation [20]. Additionally, *BRAF*^V600E^ melanomas display basal mTORC1 activation [21, 22] and a fragmented mitochondrial network, maintained by DRP1 phosphorylation at Ser616 downstream of MAPK signalling [23, 24], linking oncogenic signalling to altered mitochondrial dynamics, metabolic rewiring and mitophagy regulation. ER stress has also been described to drive mitochondrial biogenesis and is interrelated with key signalling pathways including mTOR, AMPK, and fatty acid β-oxidation [25]. Notably, mTORC1 promotes mitochondrial biogenesis both transcriptionally, through a YY1–PGC-1α complex driving OXPHOS gene expression [26] and translationally, by enhancing the translation of nuclear-encoded mitochondrial mRNAs through 4E-BP inhibition [27]. Sustained mTORC1 activity can also initiate the UPR by increasing the protein synthesis load on the ER [25], establishing a potential bidirectional link between mTORC1 signalling and organelle stress in BRAF-mutant tumours. Given that mTORC1 is constitutively active in *BRAF*^V600E^ melanoma and promotes both mitochondrial biogenesis and ER stress responses, it represents a candidate mediator linking oncogenic signalling to the mitochondrial phenotype in these tumours.

We have recently shown that the channel protein connexin43 (Cx43), a transmembrane protein that forms hemichannels and gap junctions at the cell membrane, acts as a tumour suppressor in BRAF-mutant tumours, including melanoma. Cx43 enhances BRAF/MEKi effects while alleviating drug resistance [28] through a process involving cell senescence, increased oxidative damage [29] and DNA repair pathways [28]. Cx43 is expressed ubiquitously across different cell types and is in the inner mitochondrial membrane, where it forms channel-like structures, with its C-terminus oriented toward the intermembrane space [30]. In physiological conditions, Cx43 plays an essential role in homeostasis and survival. In ventricular cardiomyocytes, Cx43 expression correlates with increased mitochondrial respiration, and its overexpression enhances complex I (CI) activity and ATP generation, while its inhibition or knockdown reduces ADP consumption, K⁺ absorption and CI function [30, 31]. Additional interactions between Cx43 and the respiratory chain proteins AIF and ETFB suggest a role in mitochondrial redox reactions [32]. Beyond mitochondria, Cx43 mediates the dissemination of deleterious signals under ER stress in liver cells, contributing to hepatic steatosis [33].

Although these studies suggest that Cx43 influences organelle homeostasis, it is unclear whether its tumour-suppressive effects in BRAF-mutated tumours are mediated through direct regulation of mitochondrial biogenesis and function, or indirectly via pathways such as ER stress. Moreover, whether mitochondrial activity modulates Cx43’s impact on BRAF/MEKi resistance remains unresolved. Here, we examined how Cx43 influences mitochondrial activity, biogenesis, and metabolic flexibility in BRAF-mutant melanoma, focusing on adaptive responses triggered by BRAF/MEK inhibition.

We show that Cx43 increases mitochondrial biogenesis and functions together with ER and mitochondrial stress responses, including activation of the ATF6 branch of the unfolded protein response (UPR). However, this programme is largely reversed by BRAF/MEKi and is therefore unlikely to underlie the enhanced therapeutic response previously attributed to Cx43. Instead, mTORC1-associated signalling and mitochondrial oxidative stress emerge as potential vulnerabilities for further investigation.

## Methods

### Cell lines and culture conditions

BRAF^V600E^ mutant cell line A375 (ATCC) was cultured in Dulbecco’s modified Eagle’s medium (DMEM) (Lonza) supplemented with 10% fetal bovine serum (FBS) (Gibco™, Thermo Fisher Scientific). Cells were maintained under sterile conditions in a laminar flow hood (Telstar BIO II A) at 37°C and 5% CO2 in a humidified incubator (SANYO CO2). To ensure experimental integrity, all cell lines were regularly screened for mycoplasma contamination and authenticated through sequencing to prevent cross-contamination. A375 cells were transfected with a vector to overexpress Cx43, as previously described [28]. Successfully transfected cells were then selected using 0.5 – 1 µg/mL puromycin dihydrochloride.

### Immunoblot analysis

Blue-native gel electrophoresis (BNGE) and in-gel activity samples were prepared as previously described [34, 35]. Cell extracts from control and Cx43-overexpressing A375 cells were loaded onto the gel for analysis of the different mitochondrial complexes. Mitochondrial proteins were extracted in phosphate-buffered saline (PBS) containing 2% dodecyl maltoside and protease inhibitors, followed by mitochondrial isolation using 200 mg/ml digitonin. The isolated mitochondria were subsequently analysed via tandem native polyacrylamide gel electrophoresis (BN-PAGE) for immunodetection of Complexes III (CIII), IV (CIV), and V (CV), as well as biochemical activity assays of Complexes I (CI) and IV (CIV). This approach enabled the study of oxidative phosphorylation (OXPHOS) complexes in their native active conformations.

Native protein samples were separated by electrophoresis using precast NativePAGE 3–12% Bis-Tris gels (Invitrogen) utilizing either 20- or 26-well configurations based on experimental requirements. Electrophoresis was conducted at a constant voltage of 70V for 17 hours at 4°C. Proteins were subsequently transferred onto polyvinylidene fluoride (PVDF) membranes using the Mini Trans-Blot transfer system (Bio-Rad) at 3 mA for 2 hours at 4°C. Membranes were incubated overnight at 4°C with primary antibodies at optimized concentrations (see Supplementary Table 1), followed by a 1-hour incubation with horseradish peroxidase (HRP)-conjugated secondary antibodies. Protein bands were visualized using X-ray films, and band quantification, alignment, and cropping were performed using Fiji software. Optical density values were normalized to β-actin for statistical analysis.

For in-gel activity assays, gel strips were loaded with 20–40 µg of mitochondrial extract. Following electrophoresis, Complex I activity was assessed using an assay buffer containing 5 mg of nitrotetrazolium blue (NTB) and 100 µl of NADH (10 mg/ml) in 10 ml of 5 mM Tris-HCl (pH 7.4) for 5 minutes. Complex IV activity was quantified following established protocols [34, 36]. Briefly, gels were incubated in a 50 mM potassium phosphate buffer (pH 7.2) at room temperature for 30 minutes. Subsequently, a reaction solution containing 50 mM potassium phosphate buffer (pH 7.2), catalase grains, 0.5 mg/ml diaminobenzidine, and 1 mg/ml cytochrome c was applied. The gels were incubated overnight and subsequently washed with the potassium phosphate buffer (pH 7.2) at room temperature to terminate the reaction. Following enzymatic activity assays, gels were scanned for densitometric quantification and analyzed using Fiji software.

### Mitochondrial isolation and subcellular fractionation

For whole-cell lysates, cells were harvested and lysed through 30-gauge syringes in lysis buffer (150 mM NaCl, 50 mM Tris-HCl pH 7.5, 5 mM EDTA pH 8, 0.5% Nonidet P-40, 0.1% SDS and 0.5% N-lauroylsarcosine) supplemented with protease inhibitors (5 µg/mL) and 1 mM PMSF. Mitochondrial fractions were isolated from cultured cells using the Thermo Scientific Mitochondria Isolation Kit for Cultured Cells (cat. no. 89874) according to the manufacturer’s instructions (Fig. 1a only). Fraction purity was confirmed by immunoblot using tubulin as a cytoplasmic marker and TOM22 as a mitochondrial marker.

**Figure 1.**
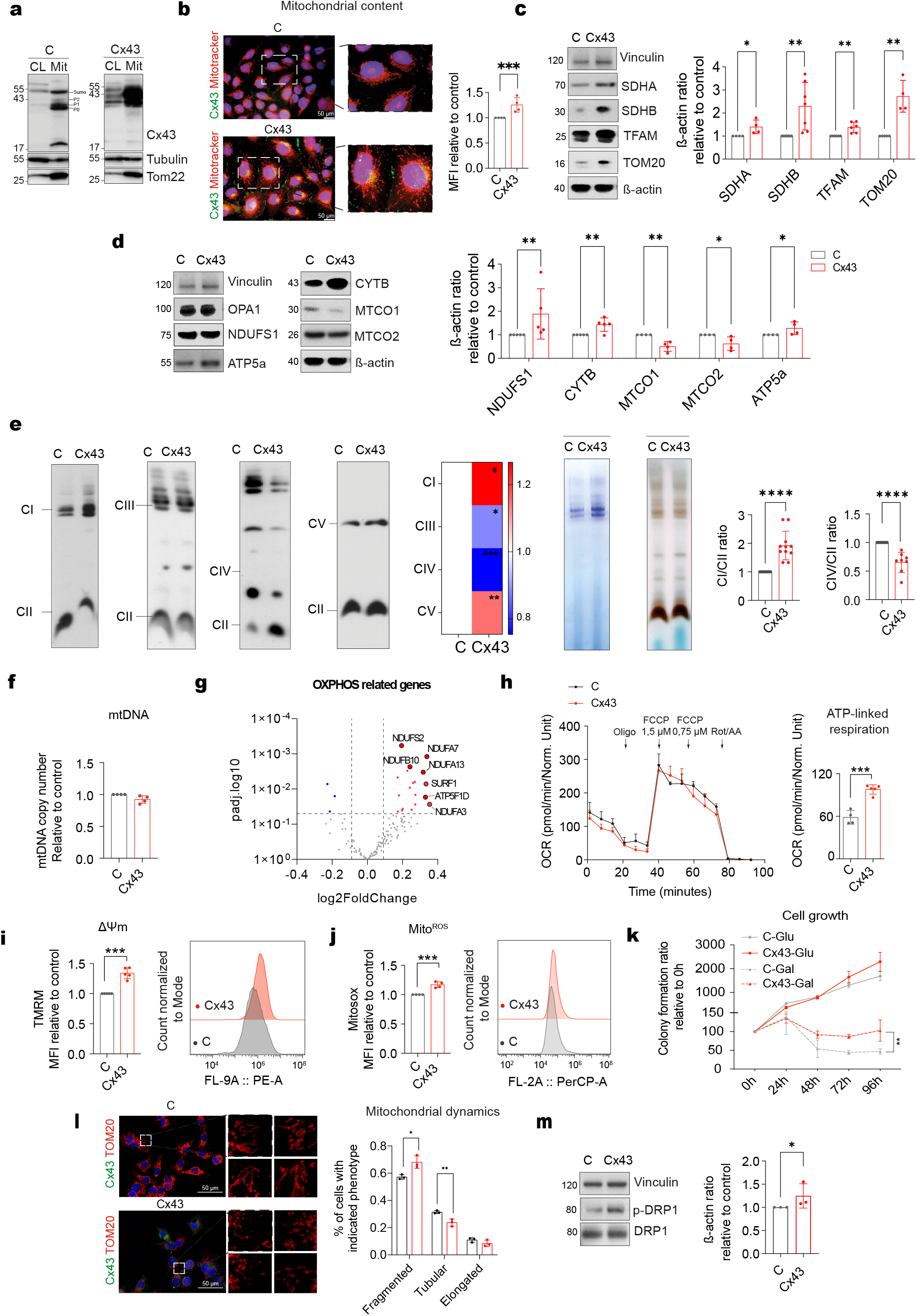
Cx43 drives mitochondrial remodelling and enhances oxidative metabolism in *BRAF*^V600E^ melanoma cells. (a) Western blot of whole-cell lysate (CL) and mitochondrial fraction (Mit) of A375-C and A375-Cx43 cells detecting Cx43. Tubulin and TOM22 were used as cytoplasmic and mitochondrial fraction markers, respectively. (b) Mitochondrial content assessed by MitoTracker (red) and Cx43 (green) immunofluorescence (left) and FACS quantification expressed as MFI (right). (c-d) Immunoblot analysis of mitochondrial biogenesis and OXPHOS subunit proteins in A375-C and A375-Cx43 cells. β-actin or vinculin were used as loading controls. (e) Immunoblots from BN and *in-gel* activity assays (right) in A375-C and A375-Cx43 cells. Heatmap of relative band intensities as well as CI and CIV activity are shown corrected by CII. (f) Relative mtDNA copy number in A375-C and A375-Cx43 cells. (g) Volcano plot of OXPHOS genes (MitoCarta3.0, n = 139) in A375-Cx43 versus A375-C cells (n = 4 per group, DESeq2 Wald test). Genes with nominal p < 0.05 are highlighted in red (increased) or blue (decreased). Only NDUFS2 and NDUFA7 remain significant after Benjamini-Hochberg correction (adjusted p < 0.05). (h) OCR following sequential injection of oligomycin, FCCP (1.5 and 0.75 µM) and rotenone/antimycin A (Rot/AA), with ATP-linked respiration quantification on the right. (i) MIMP measurement expressed as MFI relative to untreated control. (j) Mitochondrial ROS levels expressed as MFI relative to untreated control. (k) Proliferation rate of A375-C and A375-Cx43 cells over 96 hours in glucose- or galactose-enriched media, expressed as colony formation ratio relative to 0h. (l) Representative immunofluorescence images of TOM20 (red) and Cx43 (green) with quantification of mitochondrial morphology phenotypes (fragmented, tubular, elongated). (m) Immunoblot detection of p-DRP1Ser616 and total DRP1 in A375-C and A375-Cx43 cells. Vinculin was used as loading control. All experiments were performed in a minimum n=3 independent biological replicates. Data are expressed as mean ± SD. Two-tailed Student’s *t*-test and one-way ANOVA: \**P* < 0.05, \*\**P* < 0.01, \*\*\*\**P* < 0.0001.

### Immunofluorescence

Cell immunofluorescence was performed as previously described ^24^. Briefly, the cells were cultured on coverslips and subsequently fixed with 4% paraformaldehyde for 15 minutes at RT. Following a 30-minute incubation at RT, the samples were washed and blocked with a 1% BSA solution in PBS for 30 min. Subsequently, the fixed and permeabilized cells were incubated with an anti-Cx43 antibody (Supplementary Table 1)(1:200 dilution) and 20 nM MitoTracker (Thermo) or anti-TOM20 antibody (Supplementary Table 1)(1:1000 dilution) for visualizing the mitochondria. Nuclear staining was achieved using DAPI (Sigma-Aldrich)(1:1000 dilution) for 5 min at RT. The imaging was performed using Leica TCS SP5/SP8 and Nikon A1R confocal microscopes coupled to an inverted Eclipse Ti E microscope. The subsequent image analysis was carried out using ImageJ and FIJI software platforms.

### Mitochondrial morphology analysis

Mitochondrial morphology was assessed as described previously [37] with minor modifications. Regions of interest (ROIs) of 225 µmZ were selected in the cell periphery from maximum-projection images and manually thresholded in Fiji software. Phenotypic classification was based on overall mitochondrial morphology rather than the number of free ends to facilitate the detection of subtle morphological changes across experimental groups. The mitochondrial network was classified as elongated (highly connected or linearly extended), fragmented (short, rounded, and disconnected), or tubular (intermediate morphology, neither elongated nor fragmented). The proportions of cells displaying each mitochondrial phenotype were quantified in control and Cx43-overexpressing cells under treated and untreated conditions (n ≥ 15 cells per condition).

### Flow cytometry

The data for flow cytometry analysis was obtained using a CytoFLEX LX Flow Cytometer (Becton Dickinson) with CytExpert Software. The data was analyzed using FlowJo™ Software. Alive cells were gated based on forward scatter (FSC) and side scatter (SSC) parameters to discriminate cell debris. Singlet cells were sorted from aggregates on the basis of FSC-H and FSC-A. MitoTracker™ Green FM (Thermo) at 20 nM, MitoProbe™ TMRM Assay Kit (Thermo) at 50 nM and MitoSOX™ (Thermo) at 5 μM were added to seeded cells and incubated for 30 min in darkness. Following 3 washes with PBS (5 min, x1500rpm), cells were harvested and dye incorporation was measured.

### Colony assay

Colony formation assays were performed by seeding cells at approximately 10% confluence into 96-well plates and culturing for the desired time. Treatment media were changed every 48 hours. Then, cells were fixed with 2% paraformaldehyde for 15 minutes and stained with 0.05% crystal violet. Colony images were captured using an Epson Perfection V800 photographic scanner. Quantification was performed by solubilising the crystal violet stain in 30% acetic acid for 15 minutes at room temperature and measuring absorbance at 570 nm using a NanoQuant Infinite M200 microplate reader (TECAN).

### Galactose cell growth assays

A total of 1×10⁶ cells were subcultured in a 96-well plate and allowed to grow for a period of 24 hours. Following three washes with PBS, the medium was replaced with DMEM supplemented with 10% FBS and 50 µg/mL uridine or glucose-free DMEM (Gibco, 21013-024) supplemented with 10% dialysed. FBS (Gibco, 26400-044), 25 mM galactose (Sigma Aldrich, G0750) and 1.1 mM sodium pyruvate (Gibco, 11360-070) were used. The cells were permitted to grow, and the corresponding wells were fixed and stained with crystal violet at 0, 24, 48, 72 and 96 h as previously described.

### ATP determination

ATP levels were measured in triplicate in independent experiments using the Adenosine 5′-triphosphate Bioluminescent Assay Kit (Sigma-Aldrich) according to the manufacturer’s instructions. Briefly, 5000 cells were seeded on NuncMethods 103(136102) luminescence plates and the day after the media was changed according to the experiment. On the day of the experiment, cells were incubated with 5mM 2-deoxy-d-glucose (2-DG), a synthetic analogue of glucose that causes depletion of ATP derived from glycolysis, to determine mitochondrial ATP levels. After 2 hours of incubation at 37°C, the cells were lysed and the lysates incubated with luciferin and luciferase reagents. Luminescence was then recorded using an Infinite 200PRO plate reader luminometer (TECAN). Measurements were normalised to protein content and/or viable cells.

### Lactate measurements

To measure lactate concentrations, 5000 cells were seeded on luminescence plates from Nunc (136102) in complete DMEM. After 24 h, medium was replaced by DMEM supplemented with 10 % dialyzed FBS (Gibco) in presence or absence of BRAF/MEKi (1T/100D). Metabolite concentrations were quantified using the Lactate-Glo™ Assay (Promega) after 2h, according to manufacturer’s instructions.

### mtDNA copy number

Mitochondrial gene expression was assessed as a proxy for mitochondrial content by RT-qPCR. Briefly, total RNA was isolated using TRIzol reagent (Sigma) and treated with RNase-free DNase (Invitrogen) to ensure DNA removal prior to cDNA synthesis using the SuperScript® VILO™ cDNA Synthesis Kit (Invitrogen). RT-qPCR was performed using LightCycler 480 SYBR Green I Master on a LightCycler® 480 System (Roche). Expression of the mitochondrial gene *MT-ND1* was normalised to the nuclear reference gene *B2M* and used as a measure of relative mitochondrial transcript levels.

### Transcriptomic analysis

RNA sequencing data used in this study were previously generated and reported in Varela-Vázquez & Guitián-Caamaño et al. (2025) [28] and are publicly available through the NCBI Sequence Read Archive (SRA) under BioProject accession number <u>PRJNA1225446</u>. Briefly, RNA-seq was performed on A375 BRAF^V600E^ mutant melanoma cells overexpressing Cx43 or transfected with an empty vector control (A375-C), with four biological replicates per condition, after 24 h in the presence or absence of BRAF/MEKi (1T/100D). Only untreated cells were analysed in the present study. Differential expression between A375-Cx43 and A375-C cells was tested with the Wald test in DESeq2 (v1.38.3), as described in [28], and log2 fold changes are reported without shrinkage. From these results, we examined individual genes related to OXPHOS, as defined by the MitoCarta3.0 MitoPathways database, and a curated set of genes related to cellular stress responses, covering the ER and mitochondrial unfolded protein responses (UPR), general stress, mTOR signalling and one-carbon metabolism. Genes with nominal p < 0.05 are highlighted in the figures, and those that remain significant after Benjamini-Hochberg correction (adjusted p < 0.05) are indicated in the figure legends.

### Oxygen consumption rate and extracellular acidification

Oxygen consumption rate (OCR) and extracellular acidification rate (ECAR) were determined using a Seahorse XF96 extracellular flux analyzer and XF Cell Mito Stress Test Assay (Bioscience). 17k cells were seeded onto XF96 microplates in complete DMEM, in presence or absence of BRAF/MEKi for 72 h. Then, the medium was replaced by fresh media without H2CO3 (#M0286, Sigma) supplemented with 4.5 g/l glucose, 3.9 mM glutamine and 1mM pyruvate 1h before analysis. Selective inhibitors were injected during analysis: oligomycin (1 μM), carbonyl cyanide 4-trifluoromethoxy phe-nylhydrazone (FCCP; 0.75 and 1.5 μM) and rotenone/antimycin A (1 μM). Data were normalized for total protein content using the BCA Protein Assay Kit (Thermo Scientific). OCR trace measurements were calculated following previously described specifications [38].

### Statistical analysis

Unless otherwise stated, statistical analysis was performed with GraphPad Prism 9.4.1 software. Data is represented as mean ± SD. Differences between two groups were estimated with two-tailed unpaired Student’s t test or the Mann–Whitney U-test. Statistical significance was determined using two-way ANOVA followed by Holm–Šídák post-hoc for experiments with multiple variables. A minimum sample size of 3 independent experiments was used in all experiments for statistical comparisons. P value < 0.05 was considered statistically significant. *P< 0.05, **P< 0.01, ***P< 0.001, ****P < 0.0001.

## Results

### Cx43 modulates OXPHOS dependent metabolic rewiring and mitochondrial dynamics in BRAF-Mutant Melanoma Cells

We recently demonstrated that Cx43 expression exerts anti-tumour activity in *BRAF*^V600E^ mutant tumours and enhances the effect of BRAF/MEKi [28]. Since Cx43 is known to modulate mitochondrial CI activity in cardiomyocytes [30] and mitochondrial transport in osteocytes [39], we wondered whether its tumour-suppressive effects are linked to metabolic rewiring. To directly evaluate the effect of Cx43 on mitochondrial biogenesis and function we made use of our previously established model of Cx43-expressing *BRAF*^V600E^ mutant melanoma cells [28] (**Fig.1a**). Subcellular fractionation confirmed the presence of Cx43, including lower-molecular-weight isoforms, in the mitochondrial fraction of these cells (**Fig. 1a**). We observed increased mitochondrial density (**Fig.1b**), consistent with this, levels of mitochondrial biogenesis markers (SDHA, SDHB, TOM20) and the transcription factor TFAM were elevated (**Fig.1c**), along with other OXPHOS proteins (**Fig.1d**). In keeping with previous findings [30], this increase in OXPHOS proteins was accompanied by higher levels of OXPHOS CI and CV as well as increased CI *in gel* activity (**Fig. 1e**). In contrast, complexes III (CIII) and IV (CIV) were diminished, with CIV showing the greatest decrease at both free subunit levels, including the mtDNA-encoded subunits MT-CO1 and MT-CO2 (**Fig. 1e–f**). Despite these changes, the distribution of mitochondrial supercomplexes remained largely unchanged ([40]; **Fig. 1e–f**). TFAM acts by compacting mitochondrial DNA (mtDNA) into nucleoids and directly modulates mtDNA copy number [41] and is primarily regulated by PGC1a [42]. Besides increased TFAM levels (**Fig.1c**), we did not detect differences in mtDNA copy number (**Fig. 1f**), and mtDNA-encoded transcripts showed no consistent change (Fig. 1g): MT-ND2, MT-ND4 and MT-ND4L decreased and MT-ND6 increased at nominal p < 0.05, but none remained significant after correction for multiple testing. In contrast, nuclear-encoded OXPHOS transcripts showed a modest but consistent increase, with 99 of 126 genes showing a positive log2 fold change (Fig. 1g). MITF, which together with PGC1α has been associated with mitochondrial rewiring in melanoma [12, 14, 17, 43], showed a modest increase that was not significant after correction for multiple testing (log2 fold change 0.28, nominal p = 0.011, adjusted p = 0.17), whereas PGC1α (PPARGC1A) expression was too low to be assessed.

Functionally, Cx43-expressing cells displayed elevated ATP-linked respiration (**Fig. 1h**), higher mitochondrial inner membrane potential (MIMP) (**Fig. 1i**), and increased mitochondrial ROS production (**Fig. 1j**). Glycolytic activity itself was not altered, as ECAR (**Fig. S1a**) and lactate production (**Fig. S1b**) remained unchanged. To assess metabolic plasticity, cells were cultured in a galactose medium to restrict glycolysis [44] and shift metabolism towards glutamine dependence [45]. Under these conditions, Cx43-expressing cells showed better viability compared to wild-type controls (**Fig. 1k**), suggesting that Cx43 enhances metabolic flexibility by supporting mitochondrial function and cell survival when cells are forced to rely on OXPHOS.

Finally, we examined mitochondrial morphology and observed that cells overexpressing Cx43 displayed a pronounced increase in mitochondrial fragmentation (**Fig. 1l**). This phenotype was accompanied by enhanced DRP1 phosphorylation (**Fig. 1m**), consistent with an upregulation of mitochondrial fission [46]. Collectively, our results suggest that Cx43 promotes OXPHOS biogenesis and function in *BRAF*^V600E^ melanoma cells enhancing metabolic flexibility under stress.

### Stress response induced by Cx43 overactivation involves ER stress and mitochondrial UPR

ER stress has been described to drive mitochondrial biogenesis [47] and has also been linked with the control of signals through different pathways such as mTOR [25], AMPK [48, 49] or fatty acid ß-oxidation [50] (**Fig. 2a**). In addition, Cx43 localizes to the ER, where its folding and trafficking are regulated by chaperones such as ERp29 [51]. Under stress or viral infection, Cx43 is internalized into the ER–Golgi system, modulating channel function and intracellular signalling, and contributing to the intercellular propagation of ER stress responses such as the unfolded protein response [39, 52, 53]. Although previous studies have established a link between Cx43 and ER stress, the molecular mechanisms underlying ER stress–mediated upregulation of Cx43 remain poorly understood [54]. Thus, we investigated whether Cx43 upregulation–induced ER stress alters mitochondrial dynamics, or whether mitochondrial perturbations reciprocally modulate the ER and other stress response mechanisms.

**Figure 2.**
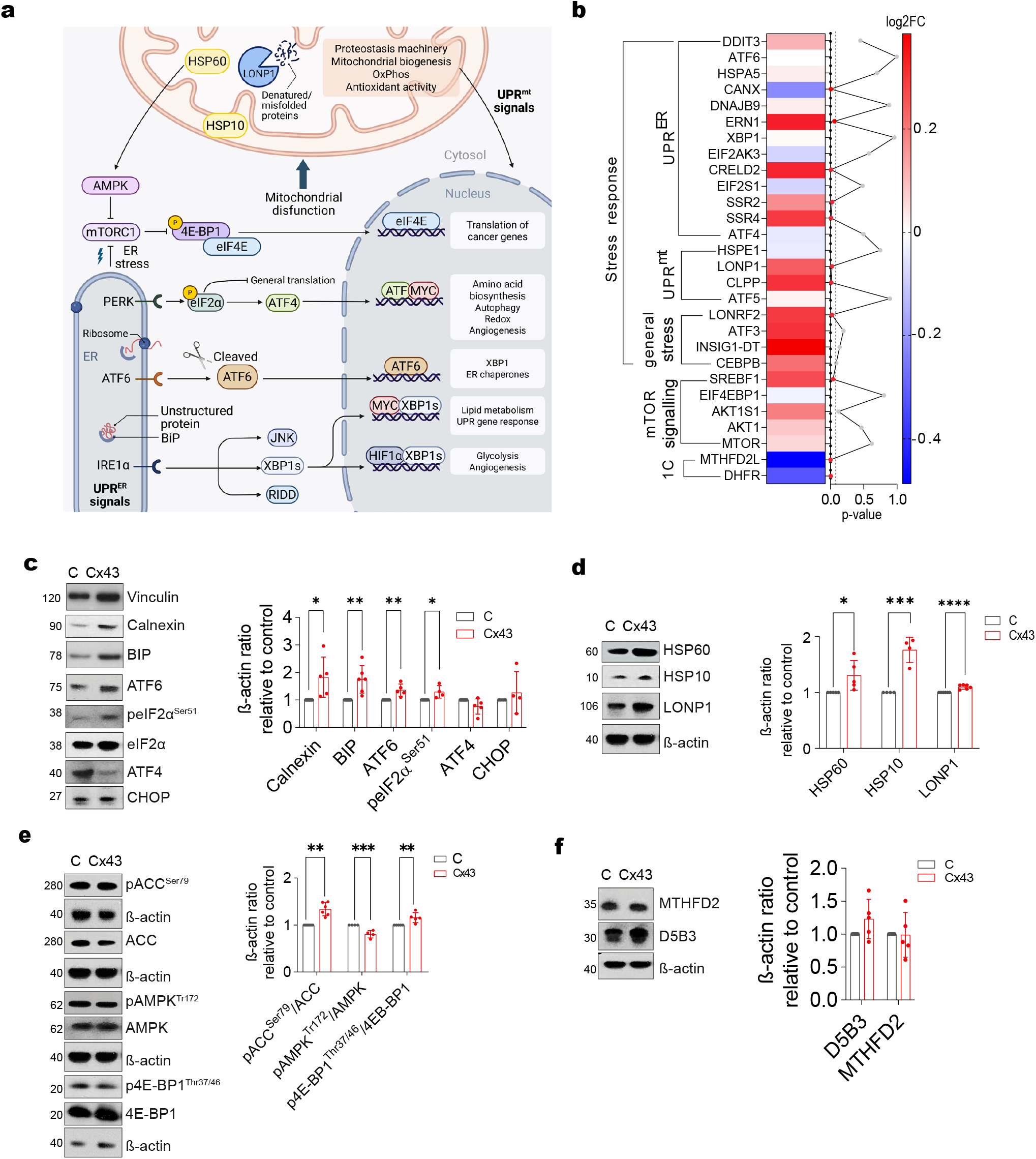
Cx43 induces ER and mitochondrial stress responses and modulates mTOR signalling in *BRAF*^V600E^ melanoma cells. (a) Schematic overview of the ER unfolded protein response (UPR^ER^) and mitochondrial UPR (UPR^mt^) signalling pathways, illustrating the three main UPR^ER^ branches (IRE1α, PERK and ATF6) and their downstream transcription factors, together with mTORC1/4E-BP1/eIF4E and AMPK/ACC signalling axes. (b) Heatmap of selected stress response and metabolic genes in A375-Cx43 versus A375-C cells, grouped by pathway category (UPR^ER^, UPR^mt^, general stress, mTOR signalling and one-carbon metabolism, 1C), showing log2FC and nominal p-value (red, p < 0.05, n = 4 per group, DESeq2 Wald test). Only MTHFD2L remains significant after Benjamini-Hochberg correction (adjusted p < 0.05). (c) Immunoblot analysis of ER stress markers (Calnexin, BIP, ATF6, peIF2α^Ser51^, eIF2α, ATF4 and CHOP) in A375-C and A375-Cx43 cells. (d) Western blot analysis of mitochondrial stress markers (HSP60, HSP10 and LONP1) in A375-C and A375-Cx43 cells. (e) Immunoblot analysis of pACC^Ser79/^ACC, pAMPK^Thr172^/AMPK and p4E-BP1^Thr37/46^/4E-BP1 phosphorylation ratios in A375-C and A375-Cx43 cells. (f) Western blot analysis of one-carbon metabolism proteins MTHFD2 and D5B3 in A375-C and A375-Cx43 cells. β-actin was used as loading control in all Immunoblot. All experiments were performed in a minimum of three independent biological replicates. Data are expressed as mean ± SD. Two-tailed Student’s *t*-test and one-way ANOVA: \**P* < 0.05, \*\**P* < 0.01, \*\*\*\**P* < 0.0001.

First, we assessed cytoplasmic and mitochondrial gene expression profiles derived from previously published RNA-seq [28] related to stress-response and metabolic pathways, such as mTOR and one-carbon metabolism (**Fig. 2a**). Transcriptomic data showed modest changes (nominal p < 0.05) in genes related to ER and mitochondrial proteostasis and to general stress responses (**Fig. 2b**). Among ER stress-related genes, Cx43 increased the expression of genes encoding proteins that support enhanced folding capacity and prolonged adaptive ER stress (**Fig. 2b**), including the calcium-binding chaperone CRELD2 and subunits of the signal sequence receptor/translocon-associated protein complex (SSR2, SSR4), a mediator of protein translocation and glycoprotein processing in the ER previously implicated in melanoma progression [55]. Increased expression of the mitochondrial proteases CLPP and LONP1 indicate parallel enhancement of the mtUPR (**Fig. 2b**). Upregulation of these genes suggests increased mitochondrial quality control at the transcriptional level in response to Cx43-induction. Beyond organelle-specific responses, a broader cellular stress was observed through the increased expression of LONRF2, an E3 ubiquitin ligase implicated in stress-response signalling and maintenance of protein homeostasis. Additionally, upregulation of the mTOR-responsive master transcription factor for lipid biosynthesis, SREBF1 (**Fig. 2b**), suggests anabolic reprogramming in the absence of increased transcription of canonical mTOR pathway components, consistent with post-transcriptional activation of mTOR signalling.

To elucidate whether transcriptional signatures affected post-transcriptional regulation, we analysed cytoplasmic and mitochondrial stress-related proteins in whole-cell lysates by immunoblotting, confirming post-transcriptional alterations (**Fig. 2c-e**). Cx43 overexpression selectively modulated UPR pathways. We observed a significant increase of the binding immunoglobulin protein (BiP), calnexin as well as activating transcription factor 6 (ATF6). This was accompanied by increased phosphorylation of eukaryotic translation initiation factor 2α (eIF2α) whereas canonical markers such as activating transcription factor 4 (ATF4) and CHOP were unchanged, indicating selective engagement of eIF2α phosphorylation and the ATF6 arm of the UPR without induction of the ATF4–CHOP axis. We also saw an increase of the mitochondrial UPR markers at protein level with upregulation of mitochondrial chaperones HSP60 and HSP10, together with an increase in the mitochondrial matrix protease LONP1 (**Fig. 2d**).

Persistent activation of mTORC1 initiates protein synthesis and UPR activation, while in the later phase induces ER stress [25]. In accordance, Cx43 overexpression resulted in a modest increase in 4E-BP1 phosphorylation (**Fig. 2e**), consistent with enhanced activity of mechanistic target of rapamycin complex 1 (mTORC1) and depression of cap-dependent translation [27, 56]. This was accompanied by reduced AMPK phosphorylation in Cx43-overexpressing cells, although phosphorylation of the AMPK substrate ACC was increased (**Fig. 2e**). In contrast, levels of MTHFD2, an mTORC1- and ATF4-regulated enzyme of one-carbon metabolism [57], were unchanged (**Fig. 2f**). Finally, the hyperfragmented mitochondrial phenotype observed in A375 cells, which is further exacerbated upon Cx43 overexpression (**Fig. 1l**), is consistent with enhanced mTOR activity. A375 cells are known to display basal mTORC1 activation relative to normal melanocytes [21, 22].

Together, these results indicate that the stress response induced by Cx43 overactivation involves mTORC1 and UPR stress responses with impact on mtUPR, potentially influencing cellular adaptation to stress conditions and driving mitochondrial biogenesis. Notably, ATF5 mRNA levels were not increased by Cx43 (**Fig. 2b**), suggesting that the adaptive ISR is probably mediated by ATF6 and ATF5-independent signalling. Overall, our data reveal that Cx43 overexpression engages both ER and mitochondrial stress responses, suggesting that Cx43 may couple ER and mitochondrial stress signalling.

### BRAF/MEK inhibition reshapes mitochondrial content, organization, and function in control and Cx43-overexpressing melanoma cells

We previously showed that Cx43 expression enhances the response of BRAF-mutant melanoma cells to combined BRAF/MEK inhibition [28]. To characterize mitochondrial responses to treatment and determine how they differ according to Cx43 expression, we exposed control and Cx43- overexpressing A375 cells to dabrafenib (100 nM) and trametinib (1 nM) for 72 h. Combined BRAF/MEK inhibition significantly reduced mitochondrial content in both cell populations, although Cx43-expressing cells retained slightly higher levels than controls following treatment (**Fig. 3a**). Several mitochondrial proteins decreased following treatment (**Fig. 3b–d**). Reductions in OPA1, NDUFS1, SDHA, CYTB, ATP5A, and TOM20 were more pronounced in Cx43-expressing cells (**Fig. 3d**).

**Figure 3.**
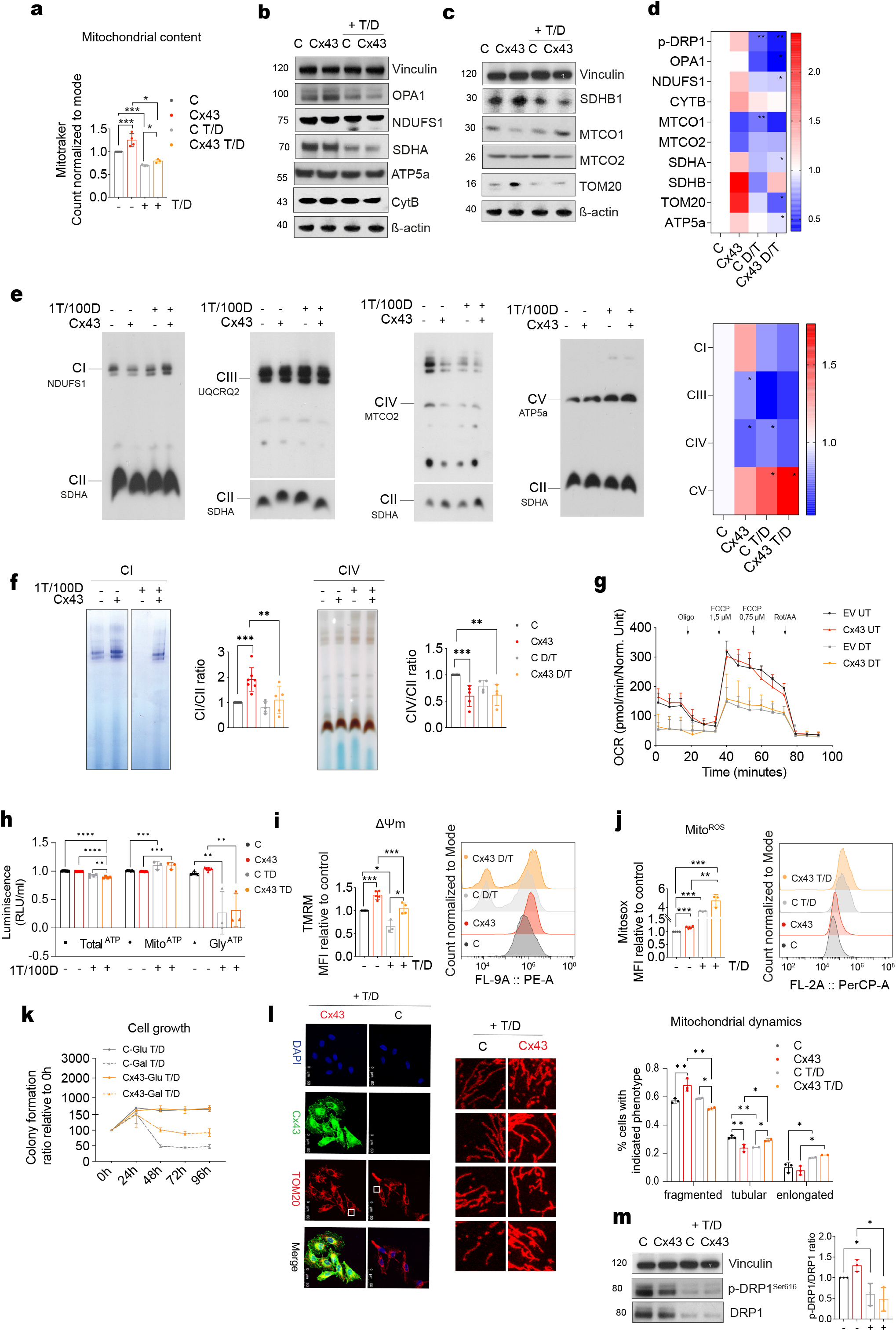
BRAF/MEK inhibition reverses the Cx43-associated mitochondrial phenotype and imposes oxidative stress. (a) Mitochondrial content in A375-C and A375-Cx43 cells treated with BRAF/MEKi (1nM T/ 100 nM D) for 72h. (b-c) Immunoblot analysis of mitochondrial OXPHOS subunit proteins in A375-C and A375-Cx43 cells ± T/D for 72h. β-actin and vinculin were used as loading controls. (d) Heatmap of mitochondrial protein levels across conditions (C, Cx43, C T/D, Cx43 T/D). (e) BN assessing ETC complex assembly in A375-C and A375-Cx43 cells ± T/D for 72h. Heatmap of relative complex band intensities across conditions is shown. (f) *In-gel* activity assays for CI and CIV with CI/CII and CIV/CII activity ratios. (g) OCR following sequential injection of oligomycin, FCCP (1.5 and 0.75 µM) and rotenone/antimycin A in A375-C and A375-Cx43 cells ± T/D for 72h. (h) Total, mitochondrial and glycolytic ATP levels in A375-C and A375-Cx43 cells ± D/T, expressed as luminescence (RLU/ml) relative to untreated control. (i) MIMP expressed as MFI relative to untreated control. (j) Mitochondrial ROS expressed as MFI relative to untreated control. (k) Proliferation rate of A375-C and A375-Cx43 cells over 96 hours in glucose-or galactose-enriched media in the presence of T/D, expressed as colony formation ratio relative to 0h. (l) Representative immunofluorescence images of Cx43 (green) and TOM20 (red) in A375-C and A375-Cx43 cells ± T/D, with quantification of mitochondrial morphology phenotypes (fragmented, tubular, elongated). (m) Western blot of p-DRP1^Ser616^ and total DRP1 in A375-C and A375-Cx43 cells ± T/D. Vinculin was used as loading control. All experiments were performed in a minimum of three independent biological replicates. Data are expressed as mean ± SD. Two-tailed Student’s *t*-test and one-way ANOVA: \**P* < 0.05, \*\**P* < 0.01, \*\*\*\**P* < 0.0001.

Despite this general reduction in mitochondrial protein abundance, native complex analysis revealed increased assembled CV following treatment in both cell populations (**Fig. 3e**). CV levels were higher in Cx43-expressing cells than in controls under both untreated and treated conditions and increased further following inhibitor exposure (**Fig. 3e**). Cx43 expression also partially attenuated the treatment-associated reduction in complex I assembly. Nevertheless, complex I and IV in-gel activities decreased after treatment in both cell populations (**Fig. 3f**). Thus, differences in the abundance of selected assembled complexes were not accompanied by preservation of their measured enzymatic activities. In keeping with this, inhibitor exposure reduced mitochondrial respiration and membrane potential in both control and Cx43-expressing cells (**Fig. 3g, i**). Total ATP levels decreased slightly, and glycolytic ATP fell markedly, whereas the relative contribution of mitochondrial ATP, measured following glycolytic inhibition with 2-deoxyglucose, increased slightly in both populations (**Fig. 3h**), suggesting a relative shift towards mitochondrial ATP reliance (**Fig. 3h**). Nevertheless, detailed respiratory analysis confirmed an overall impairment of mitochondrial respiratory capacity after BRAF/MEK inhibition, with significant reductions in basal respiration, proton leak, maximal respiration, reserve capacity and ATP-linked respiration in both cell populations (**Fig. S2a**). Lactate levels remained unchanged following treatment (**Fig. S2b-c**). Although treatment reduced mitochondrial function overall, selected Cx43-associated differences persisted. Cx43-expressing cells maintained a higher MIMP under both untreated and treated conditions and exhibited higher maximal respiration than controls following BRAF/MEK inhibition (**Fig. 3g, i**). They also showed greater survival in glucose-deprived media during treatment (**Fig. 3k**). In parallel, BRAF/MEK inhibition markedly increased mitochondrial ROS, approximately three-to five-fold, with higher levels in Cx43-expressing cells than in treated controls (**Fig. 3j**).

Lastly, BRAF/MEK inhibition also altered mitochondrial morphology, producing a less fragmented network characterized by fewer, larger mitochondria in both control and Cx43-expressing cells (**Fig. 3l**). This change was accompanied by reduced DRP1 phosphorylation (**Fig. 3m**). These findings indicate that treatment-associated changes in mitochondrial organization occurred in both cell populations despite their different basal phenotypes.

Together, these results show that combined BRAF/MEK inhibition substantially reshapes mitochondrial content, organization, and respiratory function in both cell populations. Cx43 expression does not prevent these treatment-associated changes but remains associated with differences in selected respiratory complexes, membrane potential, maximal respiration, and mitochondrial ROS-associated fluorescence.

### BRAF/MEK inhibition alters metabolic signalling and organelle stress-associated proteins

Finally, to extend the characterization of the treatment response, we examined proteins associated with metabolic signalling and organelle stress. Combined BRAF/MEK inhibition increased the ratio of phosphorylated to total 4E-BP1 in both control and Cx43-expressing cells, mainly through a reduction in total 4E-BP1 with maintained phosphorylation (**Fig. 4b**). In parallel, the ratio of phosphorylated to total AMPK and ACC decreased in both cell populations (**Fig. 4b**). Treatment also reduced the abundance of several proteins associated with ER stress and mitochondrial proteostasis. These changes were particularly pronounced in Cx43-expressing cells, including reductions in ATF6, HSP60, and LONP1 (**Fig. 4a**). Thus, the increases in stress-associated proteins observed under basal Cx43 expression were attenuated following BRAF/MEK inhibition. Finally, MTHFD2 abundance decreased after treatment in both cell populations, most markedly in control cells, whereas D5B3 showed no consistent change (**Fig. 4b**). Lastly, to functionally assess mitochondrial dependence, we targeted complex I with rotenone (Rot), complex III with antimycin A (AA), and ATP synthase with oligomycin (Oligo). Under basal conditions, Cx43-expressing cells showed lower sensitivity to rotenone and oligomycin than control cells (**Fig. S3a**), consistent with their elevated expression and activity of complexes I and V. Notably, BRAF/MEK inhibition re-sensitized Cx43-expressing cells to rotenone, resulting in a selective and significant reduction in cell biomass (**Fig.S3b**). Overall, BRAF/MEK inhibition altered metabolic signalling, reduced selected organelle stress-associated proteins and modified the mitochondrial inhibitor response, particularly in Cx43-expressing cells.

**Figure 4.**
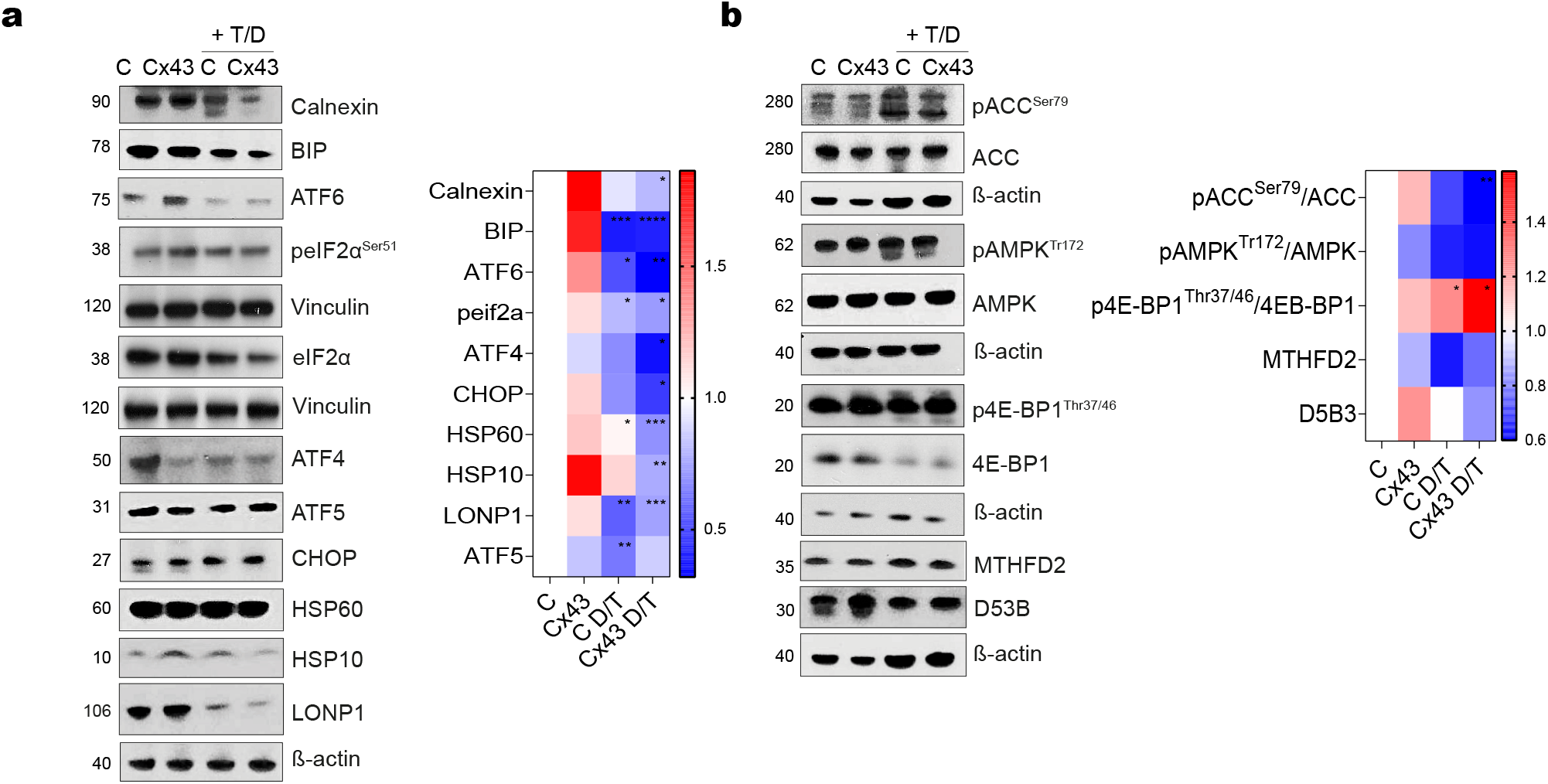
BRAF/MEK inhibition reverses Cx43-induced stress responses. (a) Immunoblot analysis of ER stress markers (Calnexin, BIP, ATF6, peIF2α^Ser51^, eIF2α, ATF4 and CHOP) and mitochondrial stress markers (ATF5, HSP60, HSP10 and LONP1) in A375-C and A375-Cx43 cells ± T/D for 72h, with heatmap of relative protein levels across conditions. Vinculin and β-actin were used as loading controls. (b) Immunoblot Analysis of pACC^Ser79^/ACC, pAMPK^Thr172^/AMPK, p4E-BP1^Thr37/46^/4E-BP1 phosphorylation ratios and one-carbon metabolism proteins MTHFD2 and D53B in A375-C and A375-Cx43 cells ± T/D for 72h, with heatmap of relative levels across conditions. β-actin was used as loading control. All experiments were performed in a minimum of three independent biological replicates. Data are expressed as mean ± SD. Two-tailed Student’s t-test and one-way ANOVA: *P < 0.05, **P < 0.01, ****P < 0.0001.

## Discussion

We investigated whether the antitumour activity of Cx43 in BRAF-mutant melanoma, including its ability to enhance the response to BRAF/MEKi [28], involves changes in mitochondrial function. Our data show that Cx43 drives a significant remodelling of mitochondria in BRAF^V600E^ cells, which is accompanied by adaptive ER and mitochondrial stress and is largely dismantled by BRAF/MEK inhibition, suggesting that the mitochondrial phenotype is largely dependent on MAPK signalling.

Cx43 increased mitochondrial content, OXPHOS mRNAs and proteins, CI assembly and activity, and CV levels, consistent with the reported role of Cx43 in supporting complex I activity and mitochondrial respiration in cardiomyocytes [30, 31, 39]. Notably, these changes occurred without consistent alterations in mtDNA-encoded transcripts or mtDNA copy number, and MITF, which together with PGC1α mediates the control of oxidative metabolism by oncogenic BRAF [12], was not significantly altered after correction for multiple testing. Instead, nuclear-encoded OXPHOS transcripts were modestly increased and the corresponding proteins were significantly upregulated, alongside TOM20 levels and 4E-BP1 phosphorylation, suggesting a combination of transcriptional and translational regulation. This remodelling was, however, unbalanced as complex IV and its mtDNA-encoded subunits MT-CO1 and MT-CO2 were decreased, while the mitochondrial proteases LONP1 and CLPP were increased. This pattern suggests an imbalance between respiratory chain components, potentially reflecting impaired synthesis or enhanced turnover of mtDNA-encoded subunits. A respiratory chain enriched in CI and CV but limited at CIV could favour electron leakage [58], consistent with increased ROS and membrane potential in Cx43 cells. Functionally, Cx43 increased ATP-linked respiration without altering glycolysis and improved survival in galactose, indicating greater oxidative competence accompanied by increased oxidative stress. This oxidative pressure may also explain why increased OXPHOS was accompanied by further fragmentation rather than fusion. BRAF^V600E^ maintains a fragmented network through ERK-dependent DRP1 Ser616 phosphorylation [59, 60], an effect further enhanced by Cx43. Mitochondrial ROS promote DRP1 Ser616 phosphorylation [61], and mTORC1 promotes fission through 4E-BP-dependent translation of MTFP1 [62]. Both mechanisms could therefore contribute to the hyper-fragmented phenotype, although their respective contributions remain to be established experimentally.

In parallel, Cx43 engaged an adaptive, non-apoptotic stress response. BiP, calnexin, ATF6 and eIF2α phosphorylation increased while ATF4, CHOP and the ATF4 target MTHFD2 were unchanged, indicating expanded folding capacity without activation of the pro-apoptotic ATF4–CHOP axis. Cx43 localisation at mitochondria-associated membranes may facilitate ER–mitochondria communication and help buffer proteotoxic stress [54, 63]. The coexistence of elevated ROS levels and increased mitochondrial fragmentation suggests that Cx43-induced stress adaptation does not fully prevent mitochondrial dysfunction. Indeed, persistent organelle stress could ultimately contribute to senescence-like phenotypes and tumour suppression, as previously reported in melanoma [64, 65] and observed in our recent work [28]. Similarly, mitochondrial stress can phosphorylate eIF2α through the OMA1–DELE1–HRI pathway [66, 67]. Increased HSP60, HSP10 and LONP1 levels, in the absence of changes in ATF5, further suggest the engagement of mitochondrial proteostasis mechanisms without detectable induction of the canonical ATF5 response. Given its reported localisation to the ER and mitochondria, and its ability to transmit ER stress between cells [33, 54], Cx43 may provide a link between ER and mitochondrial stress responses. However, direct evidence for this proposed coordination remains necessary.

Several observations are consistent with a role for mTORC1 in this programme. BRAF^V600E^ sustains constitutive mTORC1 activity through ERK/RSK in melanoma cells [68, 69], and Cx43 appeared to reinforce this anabolic state, modestly increasing 4E-BP1 phosphorylation and SREBF1 expression while reducing AMPK phosphorylation, although phosphorylation of the AMPK substrate ACC was increased. mTORC1 could contribute to each arm of the Cx43 phenotype. It promotes mitochondrial biogenesis both transcriptionally, through an mTOR-dependent YY1–PGC-1α complex that drives OXPHOS gene expression [26], and translationally, by enhancing the translation of nuclear-encoded mitochondrial mRNAs through inhibition of 4E-BPs [27]. Both routes are compatible with our data, in which modest increases in nuclear-encoded OXPHOS transcripts were accompanied by larger increases in the corresponding proteins. mTORC1 also promotes fission through 4E-BP-dependent translation of MTFP1 and restrains mitophagy [62], which would favour the accumulation of fragmented mitochondria together with increased mitochondrial mass.

The relationship between mTORC1 and organelle stress may also be bidirectional. Sustained mTORC1 signalling raises the protein synthesis load on the ER and can initiate the UPR [25], while mitochondrial dysfunction and ER stress may in turn sustain mTOR signalling through AMPK-independent mechanisms [65, 70]. Consistent with a potential link between Cx43 and this pathway, Cx43 inhibition has been associated with reduced mTOR phosphorylation in neural tissue [71]. Notably, the ATF4–MTHFD2 axis [57] was not induced, suggesting that Cx43 engages selective rather than global mTORC1 outputs. Whether mTORC1 causally underlies the Cx43 phenotype remains to be established, and testing rapamycin sensitivity and S6K signalling will be important to address this question.

BRAF/MEK inhibition reversed most Cx43-associated changes, reducing mitochondrial content, OXPHOS proteins and CI and CIV activities while promoting network elongation with reduced DRP1 Ser616 phosphorylation, consistent with previous reports [60]. The reduction in mitochondrial content contrasts with the PGC1α-dependent increase in OXPHOS described after BRAF inhibition alone [12], which may reflect the addition of MEK inhibition, the early time point analysed, or the use of an unselected population rather than drug-tolerant persisters. In contrast, Cx43 cells retained higher membrane potential, maximal respiration and ROS under treatment, but the stress proteins induced by Cx43 returned towards control levels. Notably, treatment itself increased mitochondrial ROS three- to five-fold, most markedly in Cx43-overexpressing cells, indicating that BRAF/MEK inhibition imposes a substantial oxidative burden. Because the Cx43 mitochondrial phenotype is largely MAPK-dependent, it is unlikely to explain how Cx43 enhances the response to BRAF/MEK inhibitors, which our previous work attributes to reduced DNA repair capacity and senescence [28]. The residual oxidative differences could, however, sensitise Cx43-high cells to mitochondrial inhibitors, which are active against MAPK inhibitor-resistant melanoma [72, 73]. Consistent with this possibility, Cx43-overexpressing cells showed resistance to complex I and ATP synthase inhibition under basal conditions, but increased sensitivity to complex I inhibition following BRAF/MEK treatment.

Interestingly, BRAF/MEK inhibition increased the ratio of phosphorylated to total 4E-BP1 in both cell populations, primarily by reducing total 4E-BP1 levels while phosphorylation was maintained, despite reversing most Cx43-induced mitochondrial and stress phenotypes. This is consistent with reports that BRAF inhibition can increase AKT and mTOR phosphorylation in melanoma [74], and with the established role of PI3K/AKT/mTOR activation in primary and acquired resistance to BRAF/MEKi [75, 76]. Reduced 4E-BP1 levels with maintained phosphorylation would relieve repression of eIF4E, potentially marking an early adaptive escape route, consistent with the role of eIF4F activation in resistance to BRAF and MEK inhibitors [77]. Together, these findings support further investigation of combined MAPK and mTOR inhibition and raise the question of whether Cx43 expression influences the efficacy of such combinations.

In conclusion, our results indicate that Cx43 drives mitochondrial remodelling in BRAFV^600E^ melanoma cells, characterised by increased OXPHOS biogenesis, enhanced oxidative metabolism, mitochondrial fragmentation and adaptive ER and mitochondrial stress with mTORC1 emerging as a candidate mediator. Most of these changes are attenuated by BRAF/MEK inhibition, indicating that the mitochondrial phenotype is largely MAPK-dependent and distinct from the mechanisms underlying the previously reported enhancement of treatment response by Cx43 [28]. These findings identify mTORC1 signalling and potential mitochondrial oxidative vulnerabilities warranting further investigation in the context of combination treatment strategies.

## Acknowledgements

This work was supported in part through funding from a grant from Ministerio de Ciencia, Innovación y Universidades (MICIU/AEI/10.13039/501100011033): PID2022-137027OB-I00 and PRPPID2025-168820OB-I00 ERDF/EU and a grant from HORIZON-MSCA-2023-SE-01 (EVEREST-101183034) and from EU HORIZON-CSA (TWINFLAG-101079489) (to MDM). AGD’s lab is supported by the Spanish Ministry of Science, Innovation and Universities and the State Research Agency (10.13039/501100011033) through grants PID2020-114709RA-I00, PID2023-146269OB-I00 and CNS2024-154822 as well as by a Ramón y Cajal Fellowship (RYC2020-029291-I), IDEAS222813GOME from the Spanish Association Against Cancer (AECC), and ED431F 2024/29 from Xunta de Galicia. Grant PID2023-146269OB-I00 is co-funded by the European Regional Development Fund. AG-C was funded with a predoctoral fellowship (FIS20/00310. PI: Mayán) from ISCIII. During the length of this work EF-B was funded by a predoctoral fellowship from AECC (PRDLC234093FERN. PI: Mayán and Gomez), nowadays funded by a predoctoral fellowship from the Spanish Ministry of Science, Innovation and Universities (FPU23/03704). DG-P is funded by a Xunta de Galicia Predoctoral Fellowship (ED481A-2024-165). We thank members of the CellCOM and MitoPhenomics group for helpful technical suggestions and discussions.

## Author contributions

AG-C, EF-B, CDJ-S, DG-P and PC-F designed and performed experiments. AG-C and EF-B analyzed data and prepared figures. CF-L performed the bioinformatic analysis of the RNA-seq data. TC and EF provided input, clinical advice and facilitated access to BRAF/MEK inhibitors (TAFINLAR and Mekinist). MDM and AGD conceived, directed and supervised the study. MDM, AG-C, EF-B and AGD wrote the manuscript with the input from all co-authors. All authors reviewed the manuscript.

## Data availability

RNA-seq data are available in the NCBI Sequence Read Archive under BioProject accession number PRJNA1225446 [28]. Other data that support the findings of this study are available from the corresponding authors upon reasonable request.

---

**Supplementary Figure S1.**
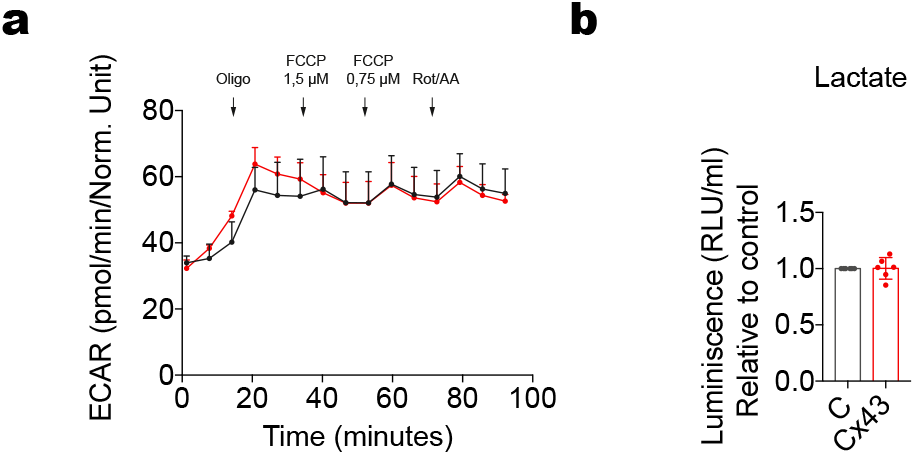
Cx43 expression does not alter glycolytic activity in *BRAF*^V600E^melanoma cells. (a) Extracellular acidification rate (ECAR) in A375-C and A375-Cx43 cells following sequential injection of oligomycin, FCCP (1.5 and 0.75 µM), and rotenone/antimycin A (Rot/AA). (b) Extracellular lactate levels in A375-C and A375-Cx43 cells, expressed as luminescence (RLU/ml) relative to untreated control. All experiments were performed in a minimum of three independent biological replicates. Data are expressed as mean ± SD.

**Supplementary Figure S2.**
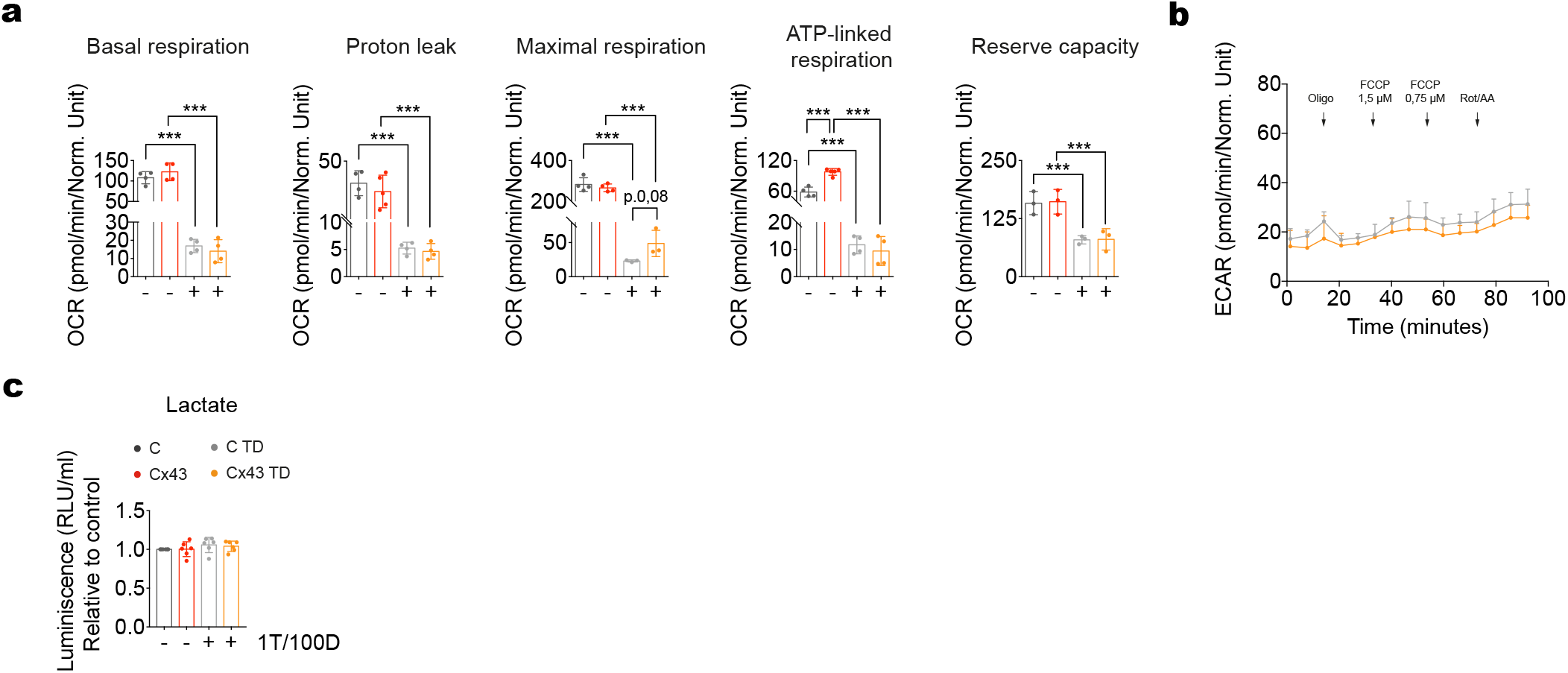
BRAF/MEK inhibition impairs mitochondrial respiration without altering glycolytic activity. (a) Basal respiration, proton leak, maximal respiration, ATP-linked respiration, and reserve capacity calculated from oxygen consumption rate (OCR) measurements in A375-C and A375-Cx43 cells treated with BRAF/MEKi (1nM T/ 100 nM D) for 72h. (b) ECAR following sequential injection of oligomycin, FCCP (1.5 and 0.75 µM), and rotenone/antimycin A (Rot/AA). (c) Extracellular lactate levels in A375-C and A375-Cx43 cells ± T/D for 72h, expressed as luminescence (RLU/ml) relative to untreated control. All experiments were performed in a minimum of three independent biological replicates. Data are expressed as mean ± SD. Two-tailed Student’s t-test and one-way ANOVA: ***P < 0.001.

**Supplementary Figure S3.**
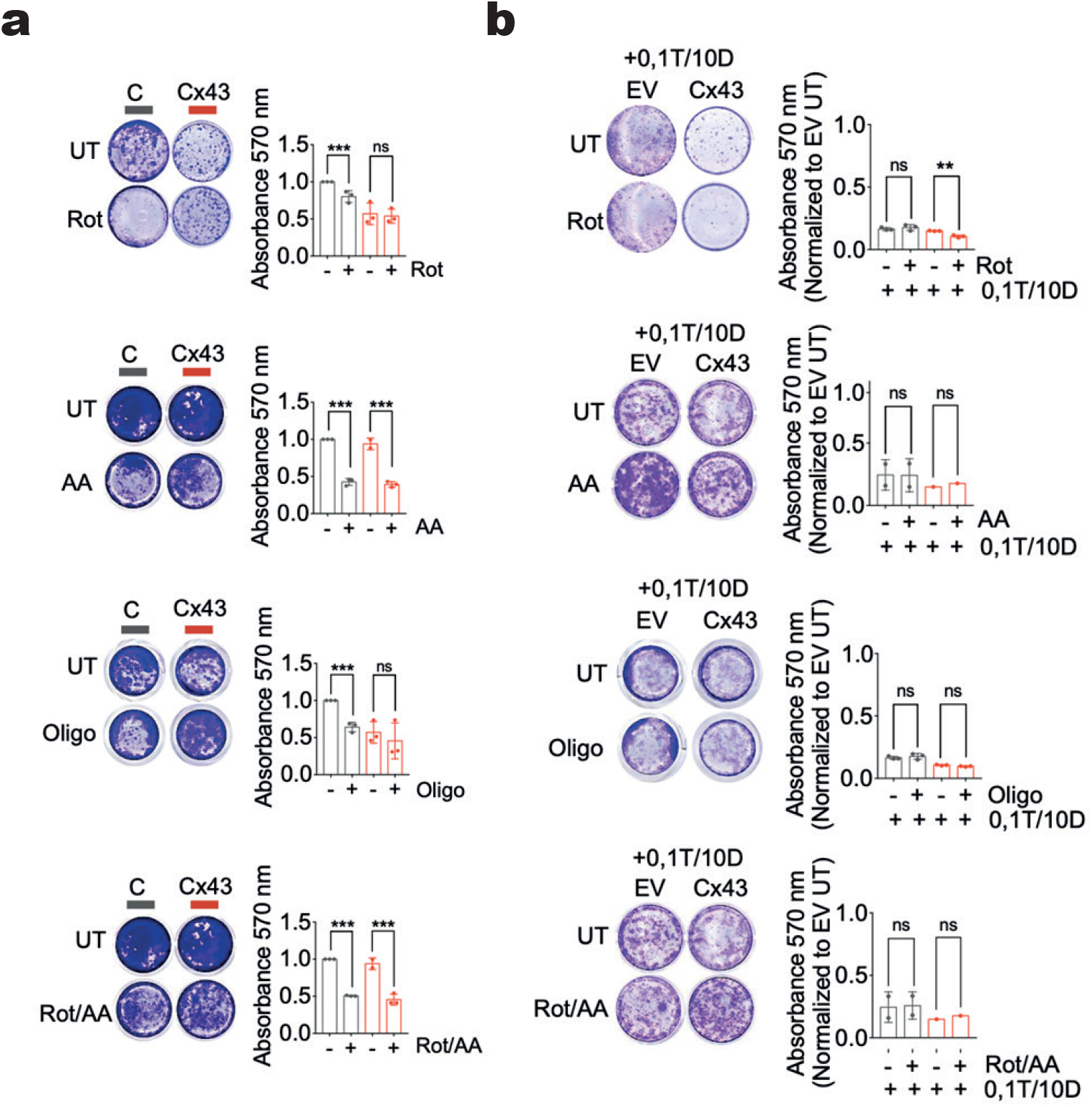
Sensitivity of A375-C and A375-Cx43 cells to mitochondrial respiratory chain inhibitors. (a) Representative crystal violet-stained colony formation assays and quantification (absorbance at 570 nm) of A375-C and A375-Cx43 cells left untreated (UT) or treated with rotenone (Rot), antimycin A (AA), oligomycin (Oligo), or rotenone/antimycin A (Rot/AA). (b) Colony formation assays of A375-C (EV, empty vector) and A375-Cx43 cells treated with the same mitochondrial inhibitors in the presence of BRAF/MEKi (0.1 nM T / 10 nM D), with quantification normalized to untreated EV cells. Data are expressed as mean ± SD. Two-tailed Student’s t-test and one-way ANOVA: **P < 0.01, ***P < 0.001.

