## Supplemental Table 1 for "Connexin43 drives mitochondrial biogenesis and organelle stress in BRAF-mutant melanoma cells"

**Table 1.** Complete list of antibodies used in this study, including working concentration/dilution, manufacturer, and catalogue number.

| | | | **Concentration** | |
|---|---|---|---|---|
| **Name** | **Distributor** | **Ref** | **WB** | **IF/IHC** |
| 4EBP1 | Abcam | ab2606 | 1:500 | |
| 53BP1 | BioTechne | 305SS | 1:500 | 1:200 |
| ACC | Cell signalling | C83B10 | 1:500 | |
| AMPK | Cell signalling | #2532 | 1:300 | |
| Anti-mouse | Dako | P0260 | 1:2000 | |
| Anti-rabbit | Dako | P0448 | 1:1000 | |
| ATF4 | Santa Cruz Biotechnology | sc-390063 | 1:300 | |
| ATF5 | Abcam | Ab184923 | 1:1000 | |
| ATF6 | Abcam | Ab37149 | 1:500 | |
| ATP5A | Abcam | AB14748 | 1:1000 | |
| BiP | Cell signalling | #3177 | 1:1000 | |
| Calnexin | Cell signalling | C5C9 | 1:500 | |
| CHOP | Cell signalling | L63F7 | 1:500 | |
| CYTB | Proteintech | 55090-1-AP | 1:500 | |
| Cx43 | Sigma-Aldrich | C6219 | 1:1000 | 1:500 |
| DRP1 | Proteintech | 12957-1-AP | 1:1000 | |
| p-DRP1 (Ser616) | Cell signalling | 3455S | 1:500 | |

| | | | | |
|---|---|---|---|---|
| eIF2α | Cell signalling | #9722 | 1:500 | |
| Goat anti-Mouse Alexa Fluor 594 | Invitrogen | A-11032 | | 1:200 |
| Goat anti-rabbit FITC-conjugated | Invitrogen | F-2765 | | 1:100 |
| HSP10 | Santa Cruz Biotechnology | sc-376313 | 1:300 | |
| HSP60 | NOVUS | NBP1-77397SS | 1:1000 | |
| LC3B | Cell Signaling | #1673868 | 1:1000 | |
| LONP1 | Abcam | Ab103809 | 1:500 | |
| MT-CO1 | Abcam | ab203912 | 1:1000 | |
| MT-CO2 | Abcam | ab110258 | 1:1000 | |
| MTHFD2 | Cell signalling | D8W9U | 1:500 | |
| NDUFS1 | Abcam | ab169540 | 1:1000 | |
| OPA1 | Abcam | Ab119685 | 1:500 | |
| phospho-ACC | Cell signalling | D7D11 | 1:500 | |
| phospho-4EBPThr37/46 | Cell signalling | #9459 | 1:500 | |
| phospho-AMPKαThr172 | Cell signalling | #2535 | 1:300 | |
| phospho-eIF2αSer51 | Cell signalling | #9721 | 1:500 | |
| pRB | Cell Signaling | #3590S | 1:1000 | |
| SDHA | Abcam | AB14715 | 1:1000 | |
| SDHB | Abcam | Ab14714 | 1:1000 | |

| | | | | |
|---|---|---|---|---|
| TFAM | NOVUS | NBP1-71648SS | 1:1000 | |
| TOMM20 | Santa Cruz Biotechnology | sc-17764 | 1:1000 | 1:500 |
| TOM22 | Proteintech | 11278-1-AP | 1:1000 | |
| UQCRC2 | Abcam | AB14745 | 1:1000 | |
| Vinculin | Sigma-Aldrich | V4505 | 1:1000 | |
| α-tubulin | Sigma-Aldrich | T9026 | 1: 10.000 | |
| β-actin | Sigma Aldrich | A1978 | 1:2000 | |
